# Assessing Aral Sea residual lake system: impact of fluctuating salinity on phytoplankton communities

**DOI:** 10.1101/2024.09.08.611860

**Authors:** Dmitry V. Malashenkov, Lajos Vörös, Aiym Duisen, Veronika Dashkova, Aidyn Abilkas, Ivan A. Vorobjev, Natasha S. Barteneva

## Abstract

The Aral Sea was once the fourth-largest inland water body in the world. However, the lake rapidly shrank over the past six decades, mainly due to the loss of inflow from one of its tributaries, the Amu Darya River. Lakes and reservoirs are traditionally characterized by static chemical and morphological parameters, leaving untouched a dynamic impact of phytoplankton changes. We used an integrated approach combining traditional microscopy and FlowCam-based imaging flow cytometry to study phytoplankton communities during the 2018 and 2019 expeditions in the Aral Sea remnant lakes system. The residual Aral Sea water bodies experienced different environmental conditions, forming hypersaline South Aral, North Aral Sea that is constantly getting freshwater, and brackish Chernyshev Bay and Tushchybas Lake with 2-8 times amplitude of salinity changes attributed to the variability in the precipitation and periodical influx of freshwater. The salinity fluctuations had an impact on the phytoplankton communities in Chernyshev Bay, making it similar to the phytoplankton of North Aral in 2018 while resembling the hypersaline South Aral phytoplankton assemblages in 2019. Multivariate analysis revealed that salinity, water temperature, ammonium, and nitrates were major contributors to explaining the variance in the sampling data. We conclude that drastic phytoplankton fluctuations occur in the two brackish water bodies in the middle of the former Aral Sea, reflecting changes in salinity.

## 1. Introduction

The tragedy of the Aral Sea is an environmental catastrophe that has developed a great concern among scientists worldwide (Micklin, 1988; Bedford, 1996; Kawabata et al., 2018; Micklin et al., 2020; Martinez-Valderrama et al., 2023). The Aral Sea has experienced a rapid decrease in water volume and reduction in water area and has segregated into several lakes. Consequently, the amount of water entering the Aral Sea decreased, leading to drastic changes in water balance, salinity, and ecosystem structure and functioning (Aladin et al., 2004; Izhitskiy et al., 2016; Micklin et al., 2020). Furthermore, despite increases in precipitation and streamflow over the entire Aral basin from 2000 to 2018, the shrinkage of the Aral Sea continues, attributed to climate change, shift in evapotranspiration (Huang et al., 2022), and increasing irrigation usage (Li et al., 2021).

Similar to the Aral Sea, many endorheic lakes worldwide are severely impacted by anthropogenic activities, leading to shrinking and increased salinization levels (Oren et al., 2010; Wurtsbaugh et al., 2017; Chaudhari et al., 2018; Bookman, 2020). For instance, Lake Urmia has experienced significant drying due to activities like dam construction and increased use of surface and groundwater for irrigation (Garousi et al., 2013; Rahimi and Breuste, 2021). Similarly, the fate of the Aral Sea remnant lakes was greatly influenced by the construction of the Kokaral dike, which resulted in the partial recovery and stabilization of the North Aral (Izhitskiy and Ayzel, 2023).

The shrinkage of the Aral Sea has had an impact on the composition of biota in the region. This was reported by the Aladin and Zavialov groups, who studied early and late stages of succession (Aladin et al., 2019; Zavialov et al., 2003; 2009; 2010; Arashkevich et al., 2009; Izhitskiy et al., 2016; Roget et al., 2017; Krupa et al., 2019; Andrulionis et al., 2022; Plotnikov et al., 2023) in conjunction with researchers in Kazakhstan and Uzbekistan. Phytoplankton, an essential part of aquatic ecosystems, responds rapidly to environmental stressors by shifts in community structure, diversity, size, and dynamics (rev. Reynolds, 2012). Phytoplankton studies of the Aral Sea during the 21st century have been infrequent and mainly focused on some specific locations (e.g., Rusakova, 1995; Kawabata et al., 1997; Mirabdullayev et al., 2004; Zhitina, 2011). Phytoplankton is comprised of an extremely diverse group of both eukaryotic and prokaryotic organisms. Different systemic grouping is much debated and used for classification, with characterization usually done by image-based taxonomy. Traditional microscopy by highly-trained individuals for decades used in phytoplankton research. However, it is labor-intensive, time-consuming and constrained by throughput with a high potential for operator-dependence and subjectivity of results (Culverhouse et al., 2003), whereas HPLC-based chemotaxonomy techniques provide averaged data at a population level identifying several distinct algal groups (Roy et al.; 2011, Swan et al., 2016; Kramer, Siegel, 2019). In the 2000s, imaging flow cytometry (IFC) was introduced for phytoplankton research (Sieracki et al., 1998; Dubelaar, Gerritzen, 2000; Sosik and Olson, 2007; rev. Dashkova et al., 2017), thereby increasing the statistical robustness of phytoplankton image analysis, and providing capabilities for images to be archived and re-examined and for generation of large image libraries and more straightforward estimation of cell size and biovolume.

Here, we aim to address the gaps in phytoplankton research in the remnant lakes of the former Aral Sea, since phytoplankton is crucial for indication of lake changes. Our approach involves using both microscopy and IFC to conduct a comprehensive and systematic analysis of phytoplankton communities of the former Aral Sea lakes along the salinity gradient from the South Aral, Uzbekistan, to the North Aral, Kazakhstan. Combining microscopy and imaging flow cytometry is an optimal approach to analyze phytoplankton diversity on a large-scale transect. This study aims to contribute to a better understanding of endorheic lakes’ fate in the changing arid environment and provide new insights into the fluctuating ecosystems of the remnant water bodies of the former Aral Sea.

## 2. Materials and methods

### 2.1. Study area

The residual water bodies of the former Aral Sea are located in a desert and semidesert region characterized by a moderate climate with strong continentality and aridity, lower and decreasing precipitation, and high seasonal and daily fluctuations in atmospheric precipitation and air temperatures (Ginzburg et al., 2010; Lioubimtseva, 2014). The increase of temperature in this region was confirmed by different research groups (Salnikov et al., 2015; Farooq et al., 2021; Bayer-Altin et al., 2024).

As of 2018-2019, the former Aral Sea was constituted of the following residual water bodies:

1. The North (=Small) Aral Sea, with a volume of around 27 km^3^, maximum depth of about 12 m, vertically mixed water column, salinity about 7–11 ppt, and an inflow from the tributary Syr Darya (Andrulionis et al., 2022);
2. The western basin of the South (=Large) Aral Sea, the deepest water body among all the residual waters of the former Aral Sea, with a maximum depth of up to 30 m, a volume of around 44–48 km^3^, thermohaline and chemically stratified water column, and a salinity about 135–140 ppt (Andrulionis et al., 2021);
3. Lake Tushchybas, a former bay of the South Aral Sea, a relatively shallow water body with a volume of 0.72–0.78 km^3^, maximum depth of 3–4 m, mixed water column, and salinity of about 60 ppt (Andrulionis et al., 2022);
4. Chernyshev Bay, a semi-isolated hypersaline part of the South Aral Sea, barely connected with it by a narrow ephemeral channel, most probably meromictic, with thermohaline and chemical stratification of water column, and with salinity varying from the surface to the bottom (maximum depth of 13–14 m) in a range of 156–236 ppt (Andrulionis et al., 2022; Izhitskiy et al., 2021). Also, there are two ephemeral water bodies, the eastern basin of the South Aral Sea and the Central Aral Sea, which seasonally reappear in cases of high water (Micklin et al., 2020) (**Suppl. Fig. S1**).

### 2.2. Field sampling and data acquisition

Samples were taken during the field expeditions to the former Aral Sea in May 2018 and May 2019. The sampling sites were located in four different aquatic areas of the former Aral Sea – the North Aral, Lake Tushchybas, Chernyshev Bay, and the South Aral (the western basin) (**Suppl. Fig. S1**). The samples were collected from the subsurface water horizon (0.5 m depth) from nearshore and open water sites (when possible). Phytoplankton samples for microscopy counting were preserved by Lugol’s iodine solution. Additionally, for the identification of phytoplankton species, qualitative samples were collected using an Apstein plankton net. Subsamples for IFC analysis were fixed with 0.5% glutaraldehyde. Physico-chemical water parameters, including water temperature, pH, dissolved oxygen (DO), conductivity, and salinity, were measured using a YSI Pro Plus multimeter (Xylem Inc., USA), CyberScan PC 300, and Eutech Salt 6+ (Eutech Instruments, Thermo Fisher Scientific, USA). Concentrations of nutrients, including nitrates, ammonium, total phosphorus (TP), and phosphates, were measured in the field using a YSI Photometer 9500 (Xylem Inc., USA) with a set of Palintest chemical kits (Palintest Ltd., UK) (**Suppl. Table S1**).

### 2.3. Phytoplankton identification and counting using microscopy

Qualitative samples were analyzed with Leica DM2500 microscope equipped with differential interference contrast (DIC) at ×400, ×630, and ×1000 magnification. Identification of phytoplankton organisms was conducted at the lowest feasible taxonomic level (mostly at the species-level). Phytoplankton samples for microscopy counting were analyzed according to Utermöhl’s method (Utermöhl, 1958). The geometric assignation technique was used to calculate biovolumes of phytoplankton cells (Hillebrand et al., 1999; Sun and Liu, 2003). Phytoplankton biovolume was converted into biomass expressed in mg·L^−1^ (Wetzel and Likens, 2000).

### 2.4. FlowCam-based IFC analysis

Environmental phytoplankton samples were examined using a FlowCam VS-4 instrument (Yokagawa Fluid Imaging Technologies, USA). The samples were recorded in laser trigger mode with the use of ×10 and ×20 objectives as described earlier (Malashenkov et al., 2021; Mirasbekov et al., 2021; Dashkova et al., 2022). Customized phytoplankton image libraries were created using VisualSpreadSheet software v. 4.0 (Yokagawa Fluid Imaging Technologies, USA), and the obtained images were classified automatically with manual inspection into different morphological groups (**Fig. 1**). For each classified group, area-based diameter (ABD) was calculated and used as a proxy in the analysis of phytoplankton size distribution.

**Fig. 1.**
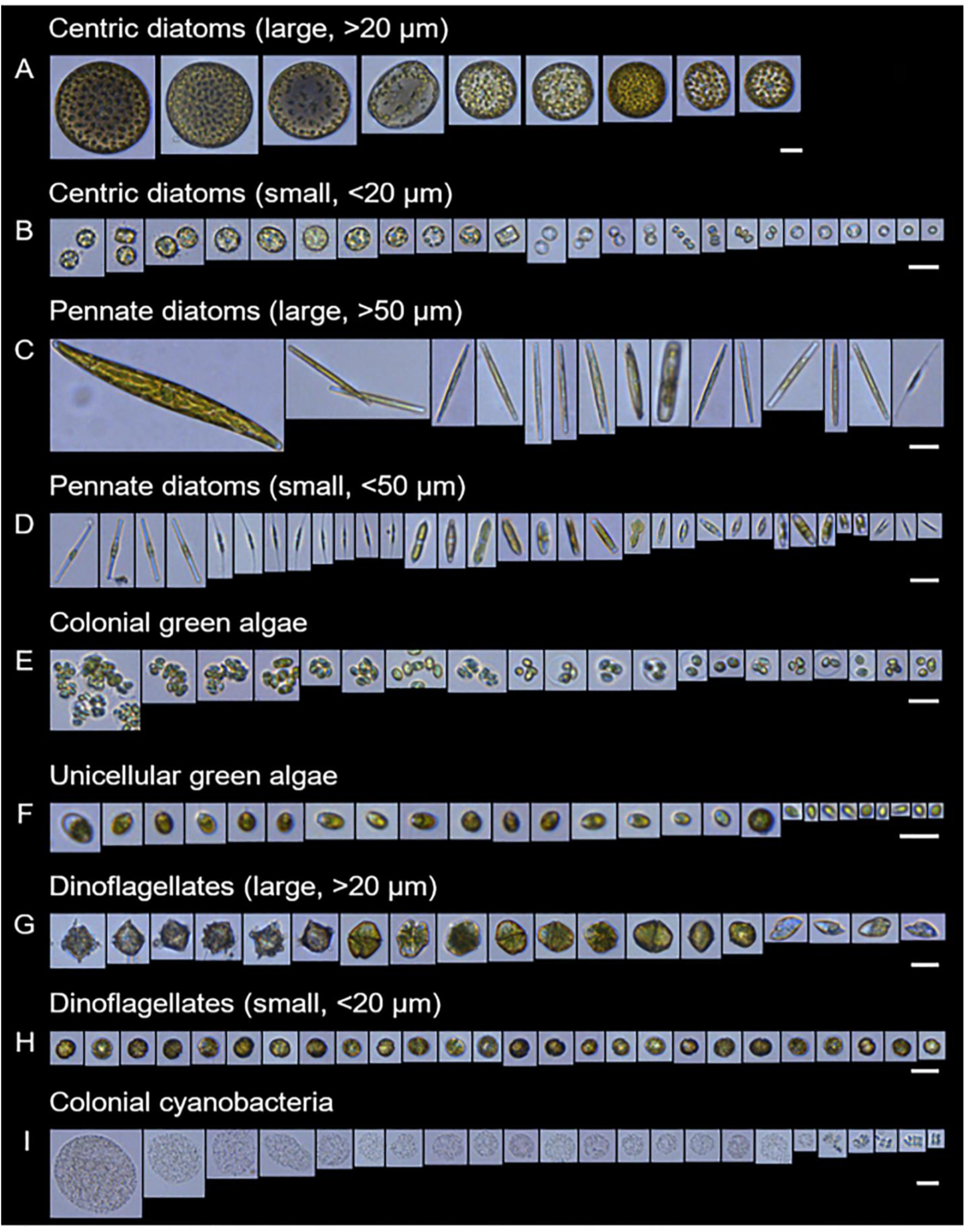
Image galleries of the FlowCam-based groups classified from the Aral Sea samples in 2018– 2019; ×10 objective, Scale bar 20 µm.

### 2.5. Data processing and statistical analysis

The identified and counted phytoplankton species were classified into functional groups (FG, coda) (Reynolds et al., 2002; Padisák et al., 2009; Salmaso et al., 2015). Diversity metrics, including Shannon (*Hʹ*) and Simpson’s (*1–D*) indices, were calculated using biomass values of each identified taxon (Magurran, 1988; Figueredo and Giani, 2001). Kruskal-Wallis test or one-way analysis of variance (ANOVA) followed by Dunn’s or Tukey’s post-hoc tests were applied to test the differences in the environmental variables and phytoplankton parameters between the seasons and the residual water bodies, depending upon the normality of variances checked with Shapiro-Wilk normality test. All calculations were done by using GraphPad Prism v. 9 (Dotmatics, USA) and PAST v. 4.02 software (Hammer et al., 2001). Spearman rank correlations between phytoplankton species were used in the construction of the species’ co-occurrence network visualized using Gephi 0.10.0 software (Bastian et al., 2009). Relationships between phytoplankton parameters and environmental variables were examined using the canonical correspondence analysis (CCA) with CANOCO 5.0 software (ter Braak and Šmilauer, 2012).

## 3. Results

### 3.1. Physico-chemical water parameters

The salinity in residual Aral water bodies was significantly different, varying from freshwater to hypersaline. In this brackish-water lake, the lowest salinity was detected for the Kokaral dam area in both years, while the highest salinity was registered at the Butakov Bay sites. On the contrary, the South Aral Sea was characterized as a hypersaline lake in both years, and salinity in Lake Tushchybas was close to marine water. In Chernyshev Bay, salinity varied from brackish to hypersaline between the years (**Suppl. Table S1**).

Considering nutrients, the Kruskal-Wallis test revealed a significant difference in nitrate concentrations among the residual water bodies (p < 0.001) but not in ammonium, total phosphorous (TP), and phosphate values. Ammonium concentration was higher in 2019 than in 2018 in the North Aral and Lake Tushchybas, while a reverse trend was observed for Chernyshev Bay and the South Aral Sea. Also, nitrate concentrations in the North Aral and Lake Tushchybas were 5-8 times higher in 2018 than in 2019 (**Suppl. Table S1**).

### 3.2. Phytoplankton species composition

A total of 233 species of nine phyla were identified in phytoplankton in the residual water bodies of the former Aral Sea basin (150 species in 2018 and 171 species in 2019) (**Suppl. Table S2**). Bacillariophyta, Cyanobacteria, and Chlorophyta were the most species-rich phyla in both years (69, 28, and 28 species in 2018 and 69, 40, 36 species in 2019, respectively). The North Aral had the highest species richness among the studied remnant Aral lakes in both years. The identified species mainly included various freshwater, freshwater/brackish and brackish diatoms, green algae, and cyanobacteria (**Fig.1**, **Suppl. Fig.2**).

**Fig. 2.**
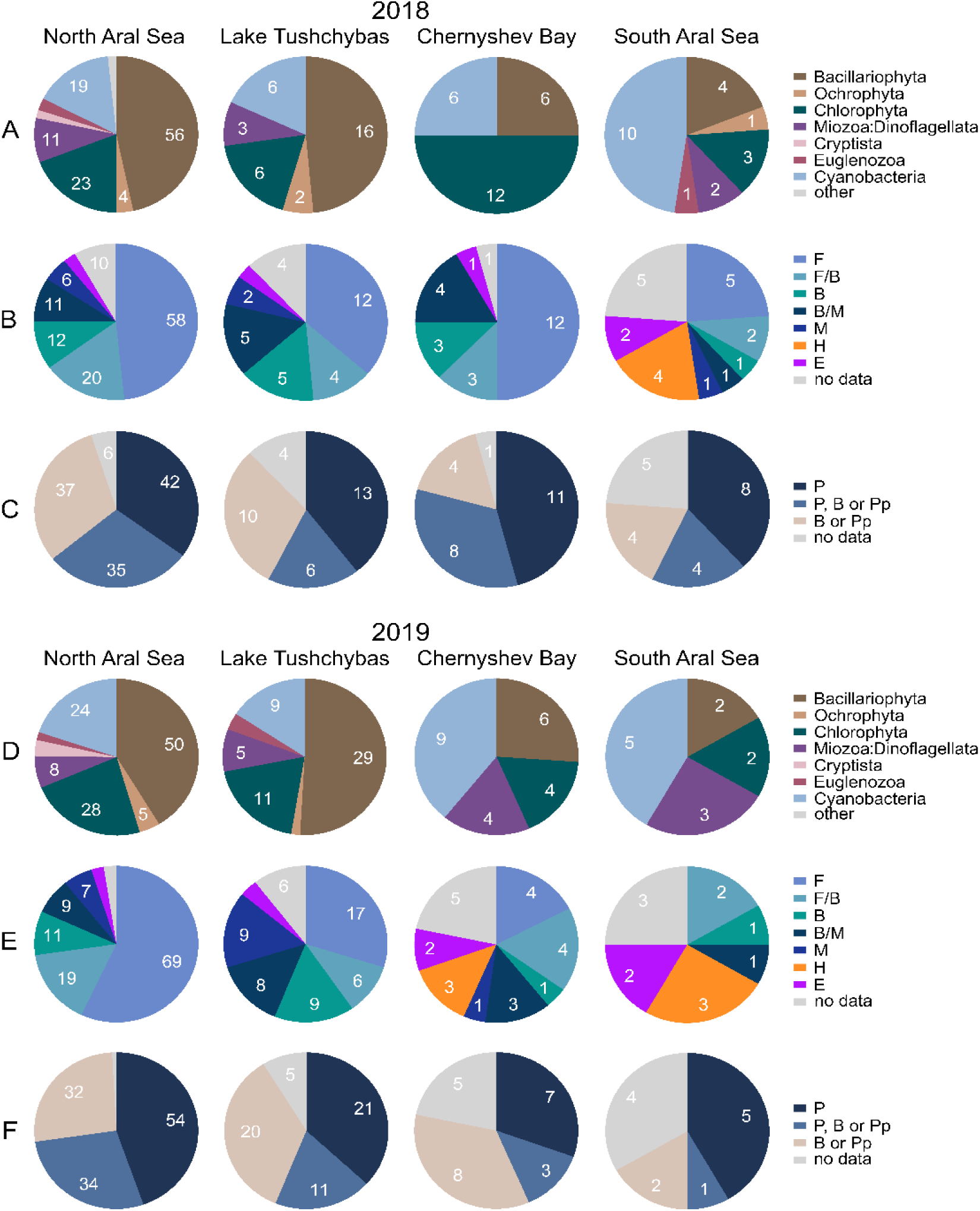
Phytoplankton community composition in the water bodies of the former Aral Sea in 2018 and 2019. **(A, D)** Contribution of different taxonomic groups (phyla) to a total number of species. **(B, E)** Contribution of species with various salinity preferences to total phytoplankton species richness: F – freshwater, F/B – freshwater/brackish, B – brackish, B/M – brackish/marine, M – marine, H – hypersaline, E – euryhaline species. **(C, F)** Contribution by species of various aquatic habitats to phytoplankton species composition: P – planktonic, P, B or Pp – planktonic-benthic or planktonic-periphytic (metaphytic), B or Pp – benthic or/and periphytic species.

By contrast, the phytoplankton composition in the South Aral, as well as in Chernyshev Bay in 2019, was a mixture of species with different salinity preferences, with a notable contribution of hypersaline and euryhaline species (**Fig. 2**). On the other hand, the community composition in 2018 featured mostly various freshwater, freshwater/brackish, or brackish species, similar to the situation in the North Aral and Lake Tushchybas (**Fig. 2**).

### 3.3. Phytoplankton biomass and biodiversity

To describe biodiversity in a quantitative way we used diversity metrics, including Shannon (*Hʹ*) and Simpson’s (*1–D*) indices. The significant difference in the diversity estimators between the residual water bodies was revealed by the one-way ANOVA (number of species, p < 0.0001; Shannon index, p < 0.01; Simpson index, p < 0.05). The North Aral stood out remarkably with a higher phytoplankton number of species and diversity, and especially against the South Aral and Chernyshev Bay (**Fig. 3A, B**; **Suppl. Fig. S3**). In the South Aral, the phytoplankton was dominated by diatoms and green algae in 2018 and colonial and unicellular picocyanobacteria (PCy) and green phytoflagellates in 2019 (**Fig. 3C**; **Fig. 4**). In Chernyshev Bay, Bacillariophyta and Chlorophyta were dominant phyla and contributed equally to total phytoplankton biomass in 2018 (**Fig. 2C**). In contrast, the colonial and unicellular PCy, diatoms, green phytomonads *Dunaliella* and dinoflagellates were dominant in Chernyshev Bay in 2019, as in case of the South Aral Sea (**Fig. 3C**; **Fig. 4**). Bacillariophyta was the most dominant phylum in the Lake Tushchybas’ phytoplankton in both years. However, the number of euryhaline species increased in 2019, whereas an assemblage of freshwater, brackish, and brackish/marine species was typical for the Lake in 2018 (**Fig. 3D**).

**Fig. 3.**
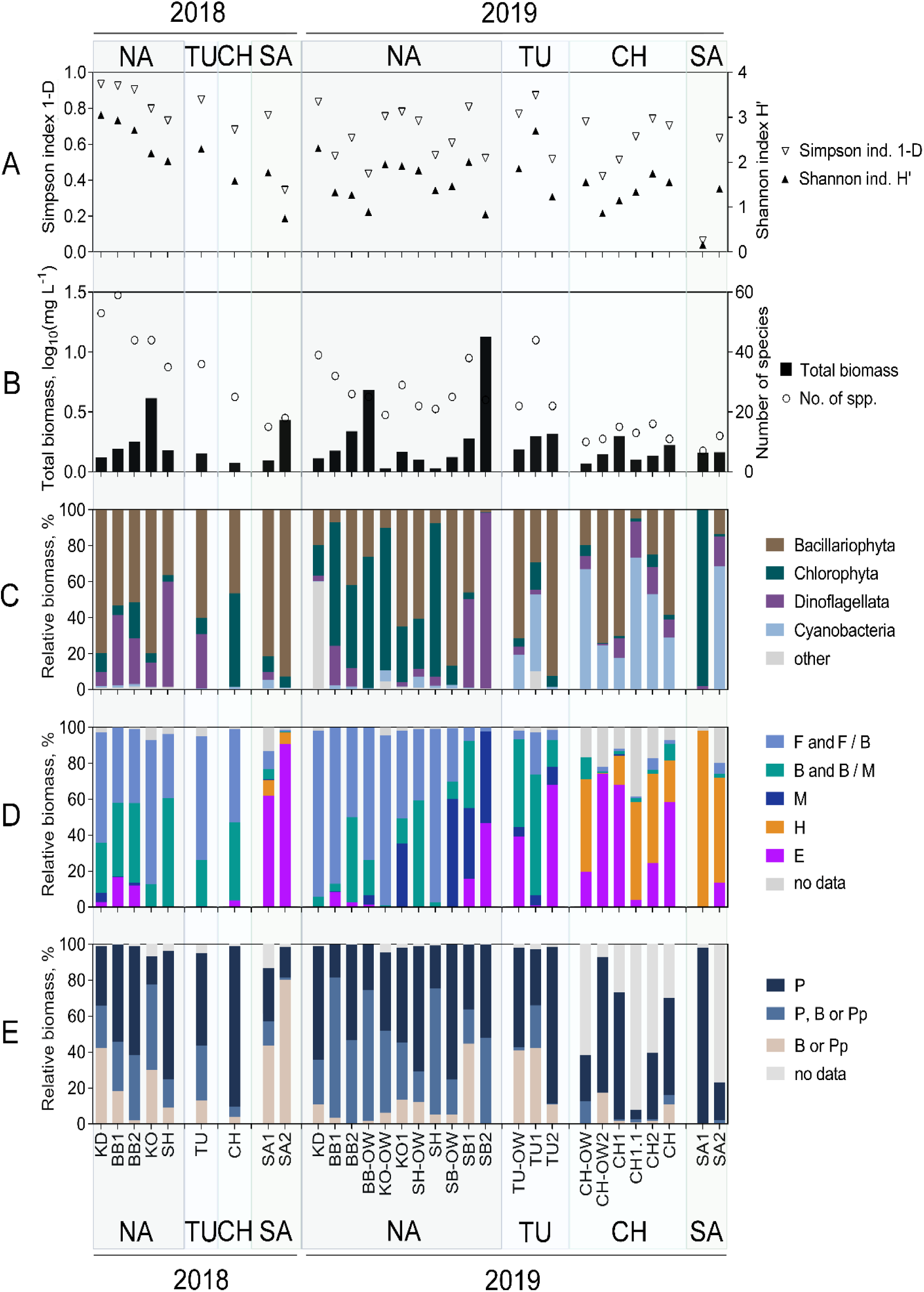
Phytoplankton total biomass, diversity estimators, and contribution of different taxonomic and ecological groups to total phytoplankton biomass in the water bodies of the former Aral Sea in 2018 and 2019. **(A)** Shannon (*Hʹ*) and Simpson (1–*D*) diversity indices. **(B)** Total phytoplankton biomass (log_10_, mg·L^−1^) and number of species. **(C)** Contribution of different taxonomic groups (phyla) to total biomass. **(D)** Contribution of species with various salinity preferences to total phytoplankton biomass: F – freshwater, F/B – freshwater/brackish, B – brackish, B/M – brackish/marine, M – marine, H – hypersaline, E – euryhaline species. **(E)** Contribution of species from various aquatic habitats to total phytoplankton biomass: P – planktonic, P, B or Pp – planktonic-benthic or planktonic-periphytic (metaphytic), B or Pp – benthic or/and periphytic species. NA – North Aral Sea, TU – Lake Tushchybas, CH – Chernyshev Bay, SA – South Aral Sea.

**Fig. 4.**
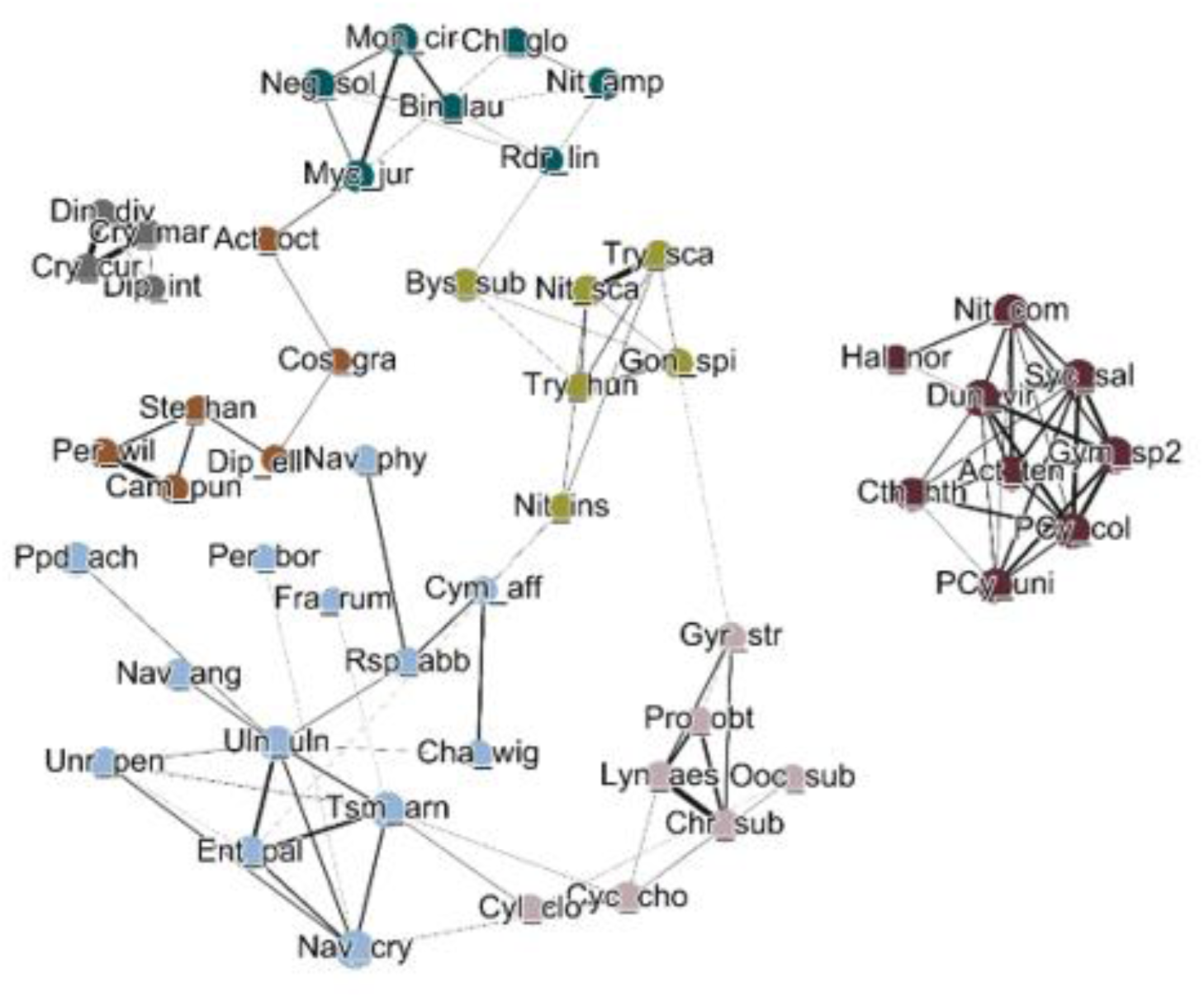
Gephi network illustrating the co-occurrence of phytoplankton species and its modular structure in the residual water bodies of the former Aral Sea. The topology of the network is based on the number of links and correlation coefficients between neighboring nodes (**Suppl. Table S3**). Each node (solid circle) represents phytoplankton species (with mean relative biomass >5%), and the size of the nodes is proportional to the number of associations. Edges (lines) indicate significant positive Spearman correlations (r > 0.5, p < 0.05), and the lines’ width corresponds to the value of the Spearman correlation coefficient between biomasses of species (correlation strength). For species codes, see **Suppl. Table S2**.

Six out of 24 functional groups (FGs or codon/coda) contributed principally to the phytoplankton biomass in the remnant water bodies of the former Aral Sea. Tychoplanktonic species of MP-codon constituted the essential part of phytoplankton biomass due to the near-shore location of most of the sites (**Suppl. Fig. S4**). The dynamics of the primarily benthic and periphytic species followed the MP-codon pattern (**Fig. 2E**). Apart from MP-codon, the assemblages of dominant FGs in the brackish North Aral were L_O_/C/D in 2018 and L_O_/C/K in 2019 (short description of dominant FGs are given in **Suppl. Fig. S4**).

Considering all the biomass data, a different degree of co-occurrences was found among the phytoplankton species. A significant nonlinear correlation (Spearman’s |r| > 0.5, p < 0.05) was found among 117 species’ pairs, which accounted for 8.8% of all of the possible pairs. The species with the greatest number of connections were *Dunaliella viridis* (8 positive and 3 negative connections) and *Actinocyclus octonarius* var. *tenellus* (6 positive and 3 negative connections) (**Suppl. Table S3; Suppl. Fig. S5**).

The ratio between unique and shared species between the different water bodies and/or years varied greatly (**Fig. 5**). A year-to-year comparison showed that the percentage of shared species was 32.4%, 23.3%, 9.3% and 43.5% of a total number of identified species in the North Aral, Lake Tushchybas, Chernyshev Bay and the South Aral Sea, respectively. In 2018, the phytoplankton community from Chernyshev Bay shared more species with phytoplankton of the North Aral. However, in 2019, the phytoplankton of the South Aral Sea shared all the identified species with the community from Chernyshev Bay (**Fig. 5**).

**Fig. 5.**
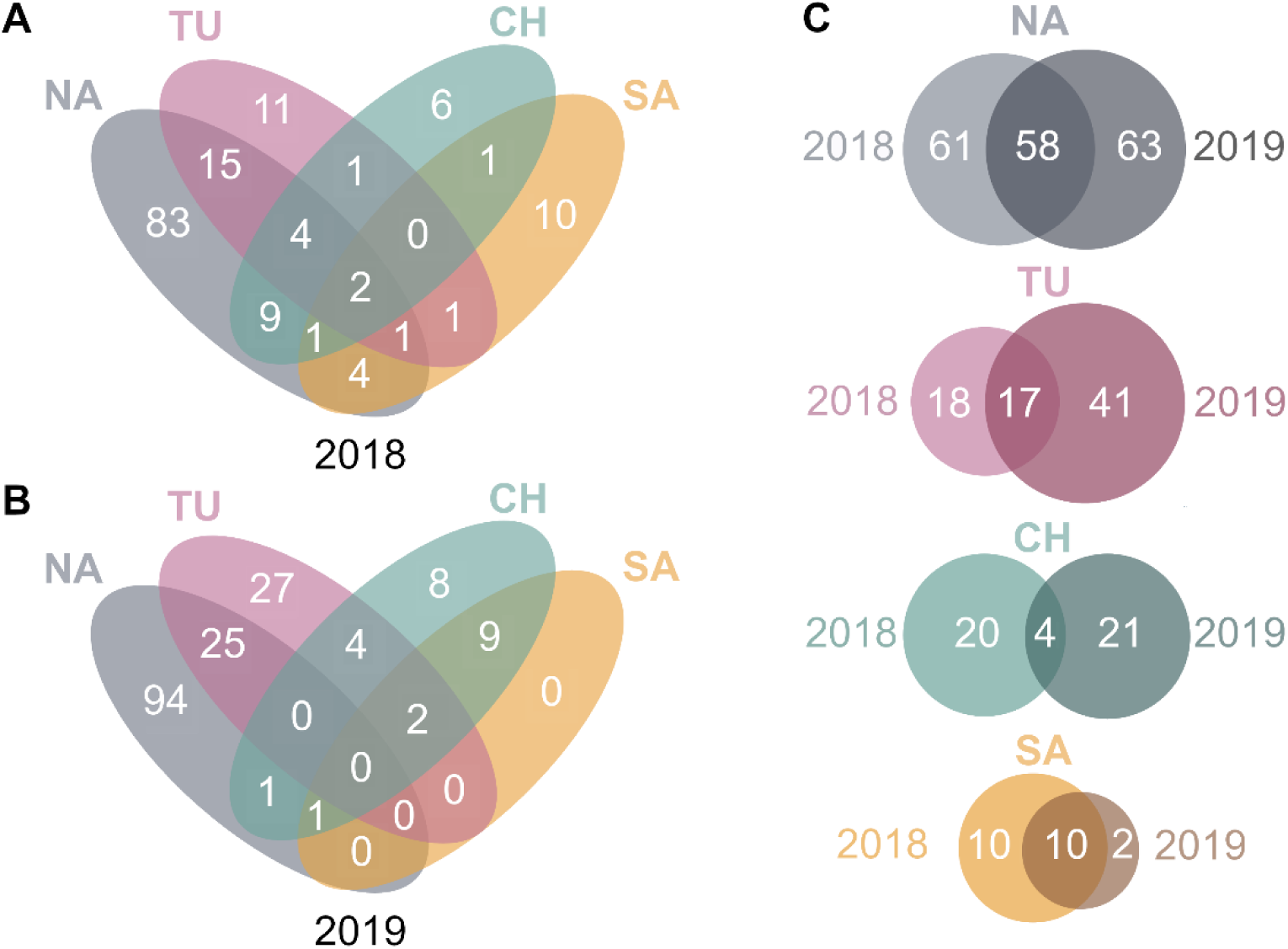
Venn diagrams of species richness across sites and years. The diagrams show the number of phytoplankton species observed between sites in 2018 (**A**) and 2019 (**B**), and a year-to-year comparison for each site (**C**). Numbers on the periphery illustrate unique species, and numbers on the intersections show shared species. Sites: NA – North Aral Sea (with 119 and 121 species in 2018 and 2019 in total, respectively), TU – Lake Tushchybas (35 species in 2018, 58 species in 2019), CH – Chernyshev Bay (24 species in 2018, 25 species in 2019), SA – South Aral Sea (20 species in 2018, 12 species in 2019).

### 3.4. FlowCam analysis of phytoplankton size distribution

The next FlowCam-based groups were distinguished: centric diatoms (small), centric diatoms algae, colonial cyanobacteria, filamentous cyanobacteria, unicellular cyanobacteria, dinoflagellates (small), dinoflagellates (large) and euglenophytes representing five taxonomic phyla (Bacillariophyta, Chlorophyta, Cyanobacteria, Miozoa and Euglenozoa). The increase in the average size of phytoplankton particles from 2018 to 2019 was noticed for the North Aral, Lake Tushchybas and Chernyshev Bay. According to the Kruskal-Wallis test, phytoplankton ABD values varied significantly for all the phytoplankton groups (p < 0.001). For several groups, significant differences in size between the sites were found. For instance, the size of large and small dinoflagellates was significantly higher in the North Aral compared to the South Aral and Chernyshev Bay in 2019 (**Fig. 6**).

**Fig. 6.**
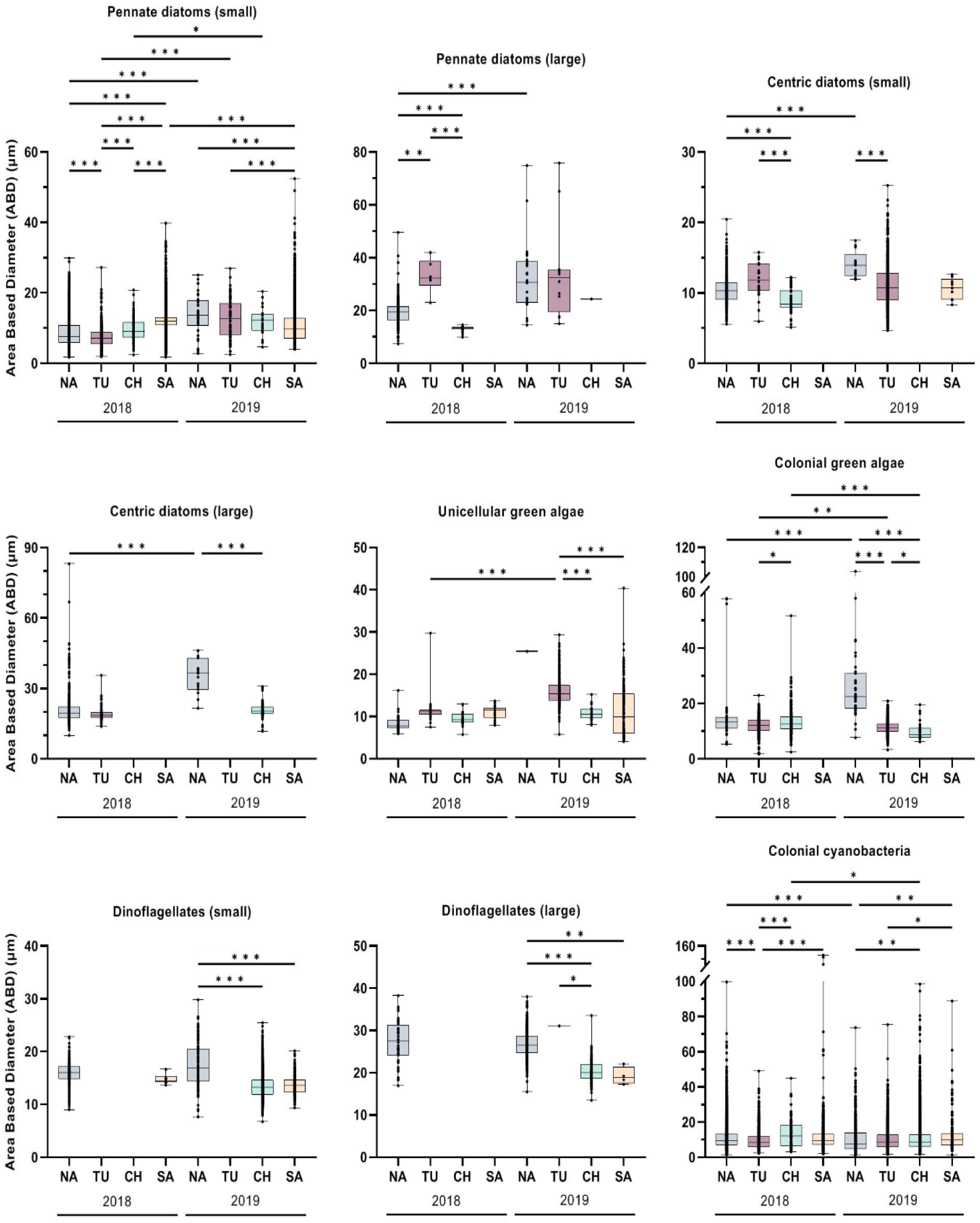
Size distribution (ABD) among the main FlowCam-based groups in phytoplankton of the Aral Sea remnant water bodies. Each boxplot shows the median, minimum and maximum values. Kruskal-Wallis and Dunn’s post hoc test: * p < 0.05, ** p < 0.01, *** p < 0.001. Water bodies: NA – North Aral; TU – Lake Tushchybas; CH – Chernyshev Bay; SA – South Aral Sea.

### 3.5. Relationship of phytoplankton community composition with environmental factors

Variation of phytoplankton species in the remnant water bodies, as well as the justification of networking-based modules of the former Aral Sea, could be mainly explained by differences in salinity, water temperature, ammonium, and nitrates according to both methods – microscopy and FlowCam IFC (**Fig. 7A, B**). Corresponding to the CCA ordination, the strongest positive relations to salinity were shown for the “module 7” species typical of the hypersaline South Aral Sea and Chernyshev Bay in 2019.

**Fig. 7.**
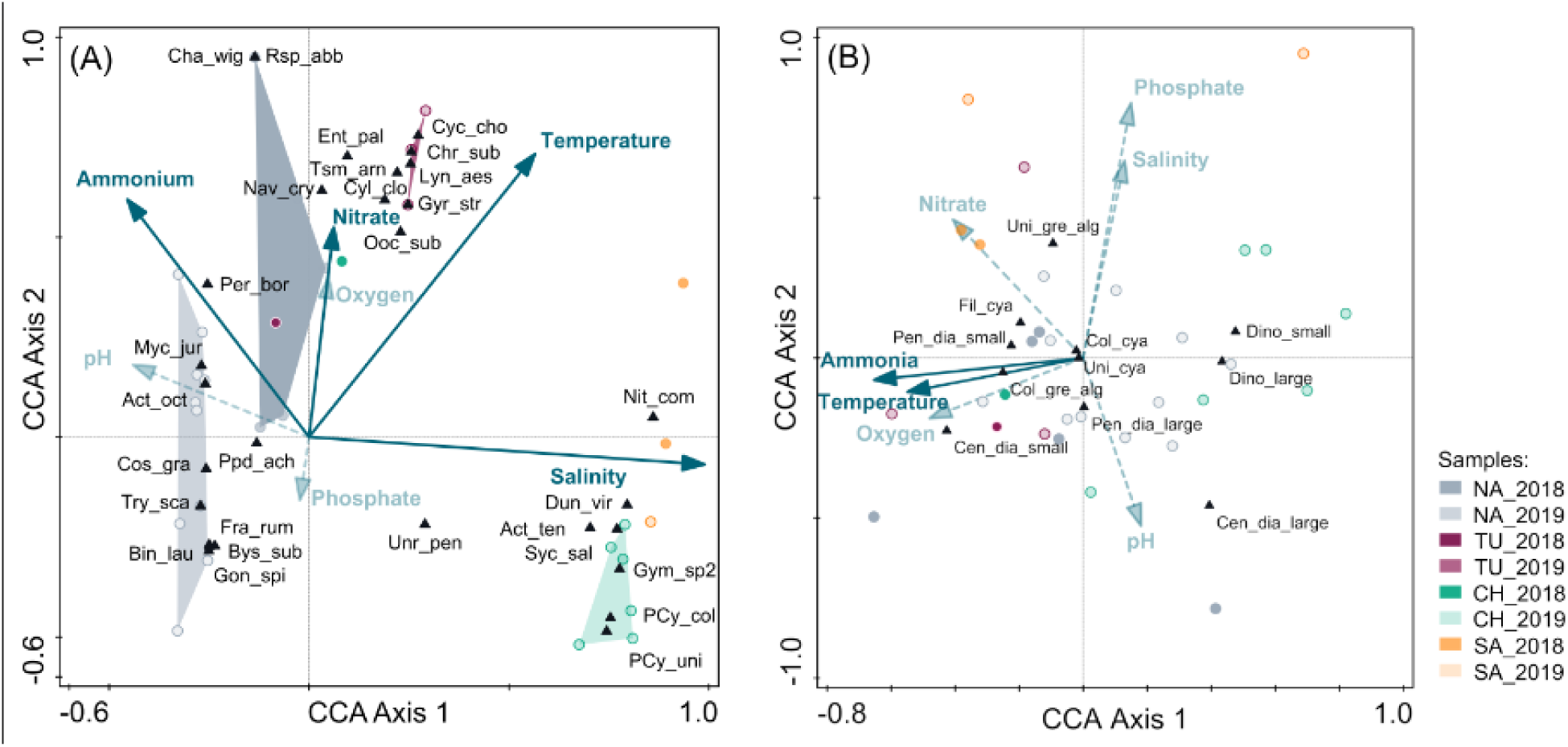
**(A)** Triplot based on canonical correspondence analysis (CCA) of the dominant phytoplankton species (with relative biomass >10%) and environmental characteristics of the residual water bodies of the former Aral Sea in 2018 and 2019. Arrows indicate quantitative environmental variables, triangles indicate phytoplankton species. Solid arrows show significant environmental parameters (Monte-Carlo permutation test, 499 permutations). All variables (except pH) were previously log-transformed. For species codes, see **Suppl. Table S2**. **(B)** CCA triplot of ABD of FlowCam-based groups and environmental variables of the former Aral Sea remnant waters in 2018 and 2019. Phytoplankton morphological group codes: Uni_gre_alg – unicellular green algae; Col_gre_alg – colonial green algae; Dino_small – dinoflagellates (small); Dino_large – dinoflagellates (large); Uni_cya – unicellular cyanobacteria; Fil_cya – filamentous cyanobacteria; Col_cya – colonial cyanobacteria; Pen_dia_small – pennate diatoms (small); Pen_dia_large – pennate diatoms (large); Cen_dia_small – centric diatoms (small); Cen_dia_large – centric diatoms (large); Eug – euglenophytes.

Regarding the functional groups (FGs), their biomass’ variations were largely explained by water temperature, nitrates, pH and salinity (**Suppl. Fig. S9; Suppl. Table S6**). Summary statistics’ output of CCA are presented in **Suppl. Tables S4–S7**.

## 4. Discussion

Phytoplankton forms a highly diverse group of prokaryotic and eukaryotic microorganisms and has been referred to as one of the paradigm systems for research on the maintenance of biodiversity (the “paradox of the plankton”) (Hutchinson, 1961; Huisman, Weissing, 1999; Stomp et al., 2004). The dynamics and structure of phytoplankton communities and primary production are affected by environmental factors such as salinity, temperature, and nutrients (Hammer, 1986; Trombetta et al., 2019; Timms, 2020; Liu et al., 2022). When salinity is low, other factors play a more significant role in community complexity and vice versa (Williams, 1998). Our data on how phytoplankton communities respond to salinity, nitrogen, and water temperature in the remnant Aral lakes aligns with this concept.

In our study, the phytoplankton assemblages distinguished by the co-occurrence network analysis were determined by the variations in the environmental variables as follows: (1) assemblage typical of the North Aral in 2018 (module 1) – tends to higher water temperature and is sensitive to nitrates; (2-3) assemblages typical of the North Aral in 2019 and from the Syr Darya mouth area in 2019 (modules 2 and 3) – influenced by increase in nitrates and ammonium; (4) North Aral assemblages (Butakov Bay 2018-2019; Shevchenko bay 2019; Koktyrnak) (module 4); (5) Shevchenko Bay’s assemblage in 2019 (module 5) – sensitive to water temperature and nitrate variations through the opposite relation; (6) assemblage typical of Lake Tushchybas in 2019 (module 6) – favoring the temperature rise; (7) species’ assemblage typical of the South Aral, and Chernyshev Bay in 2019 (module 7) – thrives in highest salinity, higher water temperature and shows strong opposite relation to ammonium. The particular interest was towards the middle part of former Aral Sea transect (Chernyshev Bay) due to the fluctuating salinity parameters highly influenced by precipitation and periodical freshwater inflow through Kokaral dike and by hypersaline water from South Aral (Barteneva et al., 2019).

We used a combination of the traditional microscopy approach and FlowCam-based imaging flow cytometry. The development of image-based analyzers (Sieracki et al., 1998; Sosik and Olson, 2007; Dubelaar, Gerritzen, 2000) permitted extensive quantification of phytoplankton. Though more taxonomic richness in the phytoplankton community structure is estimated by microscopy than measured by IFC (except for some specific algal species), the integrated approach allows for statistically robust and complementary analysis.

### 4.1 Total phytoplankton biomass

Our results show low phytoplankton biomass values in all the water bodies we studied. This finding matches the earlier studies dating back to when phytoplankton research in the Aral Sea began (Berg, 1908; Ostenfeld, 1908; Behning, 1935; Karpevich, 1960; Pichkily, 1970; Pichkily, 1981) up until the present days (Kawabata et al., 1997, 2018; Filippov et al., 2000; Klimaszyk et al., 2022). The low phytoplankton biomass and biodiversity in the South Aral are typical for hypersaline inland waters (Padisák and Naselli-Flores, 2021) and are in line with the previous studies in this area (Arashkevich et al., 2011).

### 4.2 Phytoplankton communities structure

Freshwater, brackish, and marine euryhaline diatoms, dinoflagellates, and chlorophytes dominated the North Aral, assigned to five spatiotemporal assemblages. The Kokaral dike area located near the Syr Darya River mouth, as well as semi-closed inlets such as Butakov Bay and Shevchenko Bay, represented the most stand-out areas in terms of phytoplankton results and water parameters. The increase in biomass of freshwater chlorophytes in 2019 was attributed to the freshening effect of the increased precipitation and the influence of the Syr Darya. Furthermore, the strong inflow from the Syr Darya River in 2018 charged saline Lake Tushchybas and hypersaline Chernyshev Bay with freshwater/brackish taxa from the North Aral. We conclude that Lake Tushchybas and Chernyshev Bay appear to be unstable ecosystems with high variability in environmental parameters.

The Western part of the South Aral Sea represented a typical hypersaline lake with a small number of halotolerant or euryhaline species. The study highlighted the spatial and year-to-year differences in the dominant phytoplankton taxa related to salinity increase, ammonium limitation, and possible grazing impact. In 2019, a classical phytoplankton sequence along the salinity gradient was observed. It included a large portion of halotolerant freshwater taxa in hyposaline waters (North Aral), a decrease in these species and an increase in brackish/saline/euryhaline species in moderately saline waters (Lake Tushchybas), and, finally, a restriction of community to a small number of hypersaline and euryhaline species in the hypersaline waters of the South Aral Sea and Chernyshev Bay.

### 4.3 North Aral phytoplankton

The phytoplankton in the North Aral Sea is significantly affected by nitrogen levels and water temperature. The shallow, nutrient-rich waters of the Syr Darya mouth area and the North Aral in 2019 provided a suitable environment for small-sized K-codon chlorophytes, which were the dominant species in these areas (Reynolds et al., 2002; Padisák et al., 2009).

Modern diatom flora of the North Aral was typical for lakes located in the arid regions of Central Asia and for the Aral Sea in pre-1960s era (Behning, 1935; Proshkina-Lavrenko, 1974); however, expanded in 1980s with brackish and marine euryhaline species such as *Actinocyclus octonarius* resistant to variations in salinity (Aladin et al., 2004; Aladin et al., 2009), and after the partition of the North Aral Sea remained predominant (Filippov et al., 2000). Together with planktonic species, a high contribution of tychoplanktonic diatoms (MP functional group) to phytoplankton biomass was revealed, especially at the near-shore sites. Tychoplanktonic diatoms played a significant role in phytoplankton of the North Aral in the 1990s when they dominated along with dinoflagellates (Rusakova, 1995; Orlova and Rusakova, 1999; Filippov et al., 2000). The successful survival of dinoflagellates during the years of crises lies in their salinity-tolerant nature that comes from their coastal marine origin and capacity to adapt to various mixing-irradiance-nutrient habitats (Smayda and Reynolds, 2003).

In 2019, Butakov Bay and Shevchenko Bay – the most saline sites in North Aral, had the highest contribution of dinoflagellates to the total biomass. These areas are noted for spatial heterogeneity of phytoplankton due to variations in hydrological and chemical regimes and seasonality (Koroleva, 1993; Orlova et al., 1998). Butakov and Shevchenko bays are the most remote areas from the river mouth and hence have the highest salinity, with limited water exchange with other North Aral areas, making them shelters for medium to high salt level flora and fauna (Plotnikov et al., 2017). On the contrary, the Syr Darya mouth area had predominantly freshwater and brackish colonial cyanobacteria, chlorophytes, and diatoms, some of which had been previously found downstream of the Syr Darya River (Klimaszyk et al., 2022). The increase in small-celled freshwater chlorophytes in the North Aral in 2019 was due to higher precipitation rates in the North Aral region during January-May 2019, leading to water freshening in the North Aral, except for the semi-isolated Butakov and Schevchenko bays as indicated by monitoring data from Kazhydromet (https://www.kazhydromet.kz).

### 4.4 South Aral phytoplankton assemblages

The phytoplankton assemblage of the South Aral consists of a typical hypersaline flora that is low in both biomass and diversity. However, interannual and spatial differences in dominant taxa were observed, presumably due to changes in salinity and nutrients. The South Aral phytoplankton assemblage was highly negatively related to ammonium, which is the most important nitrogen source in hypersaline waters. This is due to the sharp shifts in the uptake of ammonium along the hypersaline gradient (Joint et al., 2002). The ammonium levels in the South Aral were conditioned by the presence of the brine shrimp *Artemia parthenogenetica* (Arashkevich et al., 2009; Marden et al., 2020) and might also be linked to the high evaporation rate (Oren, 2016).

One of the keynote species, *Nitzschia communis*, is known for its ability to survive in a wide range of salinities, from freshwater to hypersaline conditions (Krammer and Lange-Bertalot, 1988; Sapozhnikov et al., 2016; Stenger-Kovács et al., 2023). Previously, it was found to be abundant in microphytobenthos of the South Aral (Sapozhnikov et al., 2010). The increase in salinity that happened in 2019 led to the development of a hard salt crust and caused a shift in the benthic community towards prokaryote dominance. At another location, *Dunaliella viridis* was predominant, possibly as a consequence of the die-off of brine shrimp due to critical temperature/salinity values (Browne and Wanigasekera, 2000).

### 4.5 Lake Tushchybas and Chernyshev Bay phytoplankton

In Lake Tushchybas, we observed a two to three-times lower salinity compared to the previous data (Aladin et al., 2009; Izhitskiy et al., 2016) due to the periodical massive inflow of fresh water from the North Aral Sea (Kazhydromet, 2020; Kazhydromet, 2021). The significant decrease in salinity of the surface waters in Chernyshev Bay in 2018, with salinity more than 8 times lower than previously recorded (Izhitskiy et al., 2016), also supports this observation. We suppose that the accumulated water mass did not mix with the pre-existing hypersaline waters, and the sharp increase in salinity the following year is likely due to extreme evaporation of surface water in the summer. This phenomenon was reflected in the phytoplankton of Chernyshev Bay, which showed a similarity to North Aral’s phytoplankton in 2018 and resembled the typical South Aral phytoplankton communities in 2019.

### 4.6 Phytoplankton cell size

Phytoplankton cell size is one of the main functional traits that can impact growth, reproduction, and survival (Litchman and Klausmeier, 2008; Zohary et al., 2017; Sonnet et al., 2022).

In the brackish North Aral Sea, the average cell and colony size of the primary phytoplankton groups increased from 2018 to 2019. This change may be attributed to the combined effects of rising salinity and falling temperatures. The decrease in water temperature in the North Aral in 2019 follows Bergmann’s rule, which suggests that phytoplankton organisms cell size diminishes along temperature gradient (Sommer et al., 2017; Cloern, 2018; Zohary et al., 2021).

The interannual variability of temperature and salinity in the hypersaline South Aral also exhibited similar patterns to those in the North Aral (salinity rose and temperature decreased in 2019), which led to an increase in the mean phytoplankton cell size. In the mesosaline Lake Tushchybas, the gain in mean cell size in 2019 could be attributed to the increase in nitrogen content, as large-cell species demonstrate a competitive advantage in nutrient-rich waters and can maintain higher rates of biomass production (Stolte et al., 1994; Cermeño et al., 2005; Hillebrand et al., 2022). While there are significant differences in cell size variability among phytoplankton groups in different Aral lakes, no clear trend has been found along the salinity gradient.

### 4.7 Large endorheic lakes shrinkage through phytoplankton glasses

The Dead Sea shares a number of similarities in history, current properties, and possible fate with the Aral Sea (Oren et al., 2010). Since 1990s, no photosynthetic plankton have been found in the lake, although previously it appeared featuring *Dunaliella* species during the dilution of the surface layer by massive winter rain floods (Gavrieli and Oren, 2004). The current state of the Dead Sea ecosystem might be interpreted as what the South Aral Sea will likely come to over time due to the ongoing salinization and water level drop (Oren et al., 2010). The ongoing desiccation crisis in one of the largest permanent hypersaline lakes, Urmia Lake, was triggered mainly by human factors such as building dams to store water for irrigation and increasing groundwater use (Pengra, 2012; Rahimi and Breuste, 2021), resulted in the loss of 90% of lake’s area (Tussupova et al., 2020). Phytoplankton of Urmia Lake was rich in *Dunaliella* spp., but following the sharp increase in salinity up to 340 ppt, the disappearance of these halotolerant microalgae is expected. The Great Salt Lake, which resembles Urmia Lake with respect to morphology, location setting, and water and sediment chemistry (Wurtsbaugh and Sima, 2022), reached its historical level minimum in 2022 primarily due to the water depletion for agriculture and other uses (Wurtsbaugh and Sima, 2022). *Dunaliella salina* survives in the North Arm of the lake (where salinity is up to 362 ppt) only when salinities decline below saturation levels (Belovsky et al., 2011). The phytoplankton shifts described in this study reflect the ongoing changes in salinity and nutrients and may serve as ecological indicators for remnant lake ecosystems.

A number of limitations of this study are important to consider. The sampling strategy and field expedition dates were complicated by remote location and limited access to the sites. Secondly, this study does not include the data on biota from different trophic levels.

## 5. Conclusions

In the present study, we used microscopy, and IFC approaches to examine the differences in phytoplankton community composition, biomass, and size distribution among the residual water bodies of the former Aral Sea in 2018-2019, as well as the environmental drivers of these changes, highlighting the unstable conditions throughout most of the Aral Sea water area.

Our findings showed that salinity, water temperature, and inorganic forms of nitrogen were the main predictors of phytoplankton biomass dynamics, dominant species composition, and phytoplanktonic cell size distribution. Additionally, we observed that the impact of temperature and nutrients based on the salinity level, affecting spatial heterogeneity and annual phytoplankton fluctuations, predominantly in the brackish waters of the former Aral Sea. The variations in salinity, temperature, and nitrogen were reflected in specific phytoplankton assemblages that were confined to a particular point in space and time, as was evidenced by the co-occurrence network analysis and confirmed by multivariate analysis. Importantly, our studies set the stage for future research on phytoplankton dynamics in the most fluctuating areas of the Aral remnant lakes.

## LIST OF ABBREVIATIONS

ABD: area-based diameter
ANOVA: analysis of variance
CCA: canonical correspondence analysis
DIC: differential interference contrast
DO: dissolved oxygen
FGs: functional groups
IFC: imaging flow cytometry
PCy: picocyanobacteria
SD: standard deviation
TP: total phosphorus

## CRediT authorship contribution statement

D.V. M. – Writing – original draft; Methodology, Data collection; Investigation, Formal analysis, Data curation, Conceptualization; L.V. - Methodology, Data collection; Investigation, Conceptualization; A.D. – Formal analysis; Data Curation; V.D. and A.A. – Methodology, Data collection; Investigation, Formal analysis; Data curation; I.A.V.- Methodology, Data collection; Investigation, Conceptualization, Supervision; N.S.B. – Methodology, Data collection; Conceptualization, Funding, Supervision, Writing - Editing. All authors – Writing – review editing.

## Acknowledgments

We would like to acknowledge the contributions of Asel Baishulakova, Kuanysh Sarkyrbayev, Aigerim Abdimanova, Polina Len, Galina Nugumanova, Adina Zhumakhanova, Kanat Samarkhanov. We are very grateful to the participants of the Aral-2018 and Aral-2019 expeditions, especially to the volunteer drivers. Our heartfelt thanks to the researchers, administration, and guides from Barsakelmes Nature Reserve for their generous help, advice, and for allowing us to use their premises for our instrumentation. Additionally, we are very grateful to QazGeo and the International Fund for saving the Aral Sea, and personally to Dr. Kanat Baigarin for his great help in organizing the expeditions.

## Appendix A. Supplementary data

The following are the Supplementary data to this article: **Supplementary Data 1**.

